# Indiscriminate activities of different Henipavirus polymerase complex proteins allow for efficient minigenome replication in hybrid systems

**DOI:** 10.1101/2024.03.18.585482

**Authors:** Xiao Li, Yanling Yang, Carolina B. López

## Abstract

The henipaviruses, including Nipah virus (NiV) and Hendra virus (HeV), are biosafety level 4 (BSL-4) zoonotic pathogens that cause severe neurological and respiratory disease in humans. To study the replication machinery of these viruses we developed robust minigenome systems that can be safely used in BSL-2 conditions. The nucleocapsid (N), phosphoprotein (P), and large protein (L) of henipaviruses are critical elements of their replication machinery and thus essential support components of the minigenome systems. Here, we tested the effects of diverse combinations of the replication support proteins on the replication capacity of the NiV and HeV minigenomes by exchanging the helper plasmids coding for these proteins among the two viruses. We demonstrate that all combinations including one or more heterologous proteins were capable of replicating both the NiV and HeV minigenomes. Sequence alignment showed identities of 92% for the N protein, 67% for P, and 87% for L. Notably, variations in amino acid residues were not concentrated in the N-P and P-L interacting regions implying that dissimilarities in amino acid composition among NiV and HeV polymerase complex proteins may not impact their interactions. The observed indiscriminate activity of NiV and HeV polymerase complex proteins is different from related viruses, which can support replication of heterologous genomes only when the whole polymerase complex belongs to the same virus. This newly observed promiscuous property of the henipavirus polymerase complex proteins could potentially be harnessed to develop universal anti-henipavirus antivirals.

**IMPORTANCE:** Given the severity of disease induced by Hendra and Nipah viruses in humans and the continuous emergence of new henipaviruses as well as henipa-like viruses, it is necessary to conduct more comprehensive investigation of the biology of henipaviruses and their interaction with the host. The replication of henipaviruses and the development of antiviral agents can be studied in systems that allow experiments to be performed under biosafety level 2 conditions. Here, we developed two robust minigenome systems for Nipah virus (NiV) and Hendra virus (HeV) that provide a convenient alternative system for studying NiV and HeV replication. Using these systems, we demonstrate that any combinations of the three polymerase complex proteins of NiV and HeV could effectively initiate the replication of both viral minigenomes, which suggest that the interaction regions of the polymerase complex proteins could be effective targets for universal and effective anti-henipavirus interventions.

## INTRODUCTION

Henipaviruses, including Nipah virus (NiV) and Hendra virus (HeV), are members of the Henipavirus genus within the *Paramyxoviridae* family. NiV and HeV represent threatening zoonotic pathogens classified as BSL-4 (biosafety level 4) agents due to their high pathogenicity and lack of available vaccines and antivirals (1, 2). Outbreaks of NiV and HeV have occurred frequently in Southeast Asia and Australia (3–6). NiV was first isolated in 1999 during an outbreak in pigs that led to subsequent cases of encephalitis among pig farmers in Malaysia and Singapore (7, 8). NiV infection can cause severe respiratory symptoms as well as fatal neurological symptoms, and the virus can spread between humans (9). HeV was identified in Australia in 1994 and is associated with severe respiratory and neurological disease in horses (10). The case fatality rates of NiV and HeV in humans is 60-100% and there are no efficacious antiviral therapeutics and licensed vaccines for human use (11, 12). To date, a vaccine to protect horses from HeV has been commercialized in Australia. The lack of equivalent prophylactics for human populations remains a critical gap in public health. Furthermore, the emergence of novel Henipaviruses such as Langya (LayV), Gamak (GAKV), and Mojiang (MojV) accentuates the dynamic landscape of this viral family, warranting heightened surveillance and the need for effective intervention strategies (13–15).

As BSL-4 pathogens, research with live NiV and HeV needs to be carried out in high containment labs imposing significant limitations to the scientific research and treatment development against these viruses. Establishment of NiV and HeV minigenome systems that can be used in the BSL-2 conditions is an effective strategy to facilitate broader research aimed at understanding the molecular mechanisms involved in virus replication, and serve as a platform for testing antivirals that target these processes (16–18). The genomes of NiV and HeV comprise a single-stranded negative-sense RNA molecule of approximately 18.2 kilobases (kb) in length. These genomes encode a repertoire of structural and non-structural proteins pivotal for viral replication and transcriptional processes (19). The nucleocapsid (N), phosphoprotein (P), and large (L) proteins are central to this machinery and collectively orchestrate the assembly of ribonucleoprotein (RNP) complexes essential for viral RNA synthesis (18, 20). In general, minigenome systems for paramyxoviruses, including henipaviruses, consist of a minigenome plasmid in which a reporter gene is flanked by the viral leader and trailer promoter sequences and three helper plasmids each expressing the N, P and L support proteins under the control of an inducible promoter. After the four plasmids are transfected into cells, the N protein coats the minigenome and minigenome replication is carried out by the viral polymerase L aided by its co-factors N and P.

Minigenome systems have been created for several *Mononegavirales.* In most cases, the helper primers need to be homologous to the minigenome parent virus or all helper plasmids need to be used as a set from the same virus that can then function with a closely related heterologous virus. For example, minigenomes for the rhabdovirus infectious hematopoietic necrosis virus can be replicated efficiently by a set of helper plasmids from the related hemorrhagic septicemia virus and vice versa. However, replication is highly inefficient and does not occur when the helper plasmids from these viruses are mixed (21). Similarly, sets of heterologous proteins worked in trans to replicate minigenomes of closely related strains of vesicular stomatitis virus, but replication did not occur when helper plasmids came from mixed strains (22). Among morbilliviruses, replication of heterologous minigenomes only happens when full sets (N, P and L) of helper plasmids are used (23). Cross-activity of polymerase and its co-factors has been also reported for the pneumoviruses human, bovine, and ovine respiratory syncytial virus (RSV) (24), however filovirus helper plasmids do not seem to work in trans even if present as a set (25). Among paramyxoviruses, it has been shown that that full-length infectious clone of Sendai virus (SeV) could be successfully rescued after co-transfection with the helper plasmid set from human parainfluenza virus 1 (HPIV1) and human parainfluenza virus 3 (HPIV3) strains, but mixing plasmids or using helper plasmids from a more distant morbillivirus or pneumovirus was ineffective (26–28).

Here, we have developed an efficient minigenome system for studying NiV and HeV replication under BSL-2 conditions as an alternative system to study virus replication. Using these systems, we found unexpected remarkable promiscuity among henipavirus polymerase complex proteins that allows efficient replication of the NiV and HeV minigenomes in hybrid systems without the need for homologous components within these viruses.

## RESULTS

### Construction of NiV and HeV minigenome systems in BSR-T7/5 cells

The henipavirus viral genomes consist of a leader sequence, 6 protein-encoding genes, and a trailer sequence (Fig. 1A). We developed minigenome systems to utilize the T7 polymerase that is constitutively expressed in BSR-T7/5 cells (29). The minigenome plasmid (MG) consists of two separate functional units (Fig. 1B). The control unit encodes an internal ribosome entry site (IRES) and an enhanced green fluorescent protein (eGFP) reporter gene under the T7 promoter that acts as a control for the minigenome transfection and for the T7 polymerase activity. The viral replicon unit includes the virus 5’ end of the L gene fragment, a reporter mCherry gene flanked by the leader and trailer sequences, and the untranslated region (UTR) of N and L genes, which are cis elements essential for viral replication and transcription. In addition, the N gene 3’ UTR sequence was added between the L gene fragment and the mCherry gene to ensure that mCherry can be transcribed. The viral replicon unit is flanked by the self-cleaving hammerhead ribozyme (Hh-Rbz) before the trailer sequence and the hepatitis delta virus ribozyme (HDV-Rbz) (30) after the leader sequence to ensure transcription products have a nucleotide length divisible by six, which is necessary for efficient replication as described by the “rule of six”.

**Fig 1.**
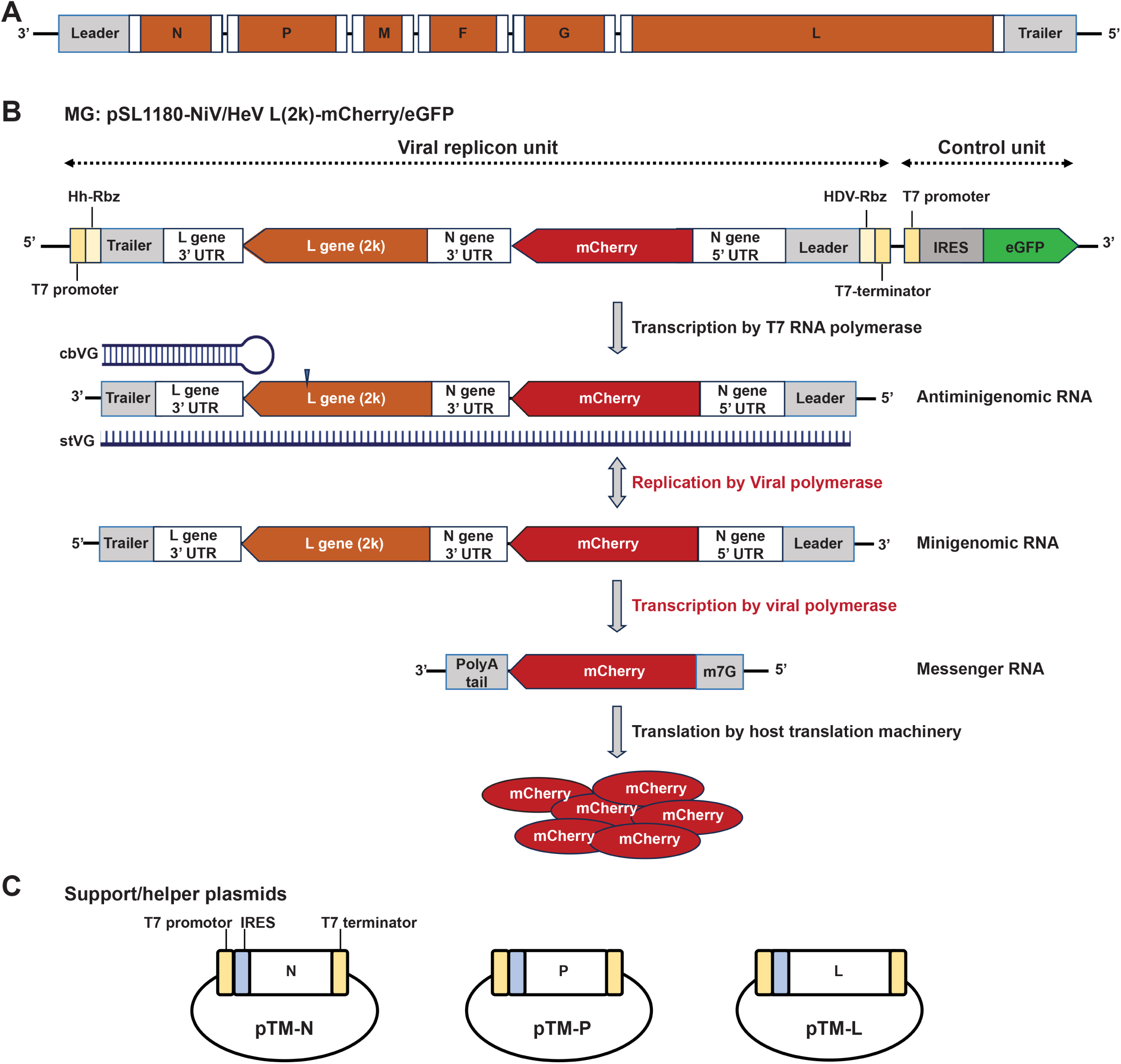
Henipavirus minigenome strategy. (**A**) Schematic representation of a generic Henipavirus genome. The six structural genes are represented by orange boxes. 3’UTR and 5’UTR of each gene are shown by white boxes. (**B**) Schematic of NiV and HeV minigenome system plasmids. RNA synthesis and translation in the viral replicon unit of minigenome system are depicted. Hh-Rbz, hammerhead ribozyme. HDV-Rbz, hepatitis delta virus ribozyme. stVG, standard viral genome. **(C)** Schematic of the three support/helper plasmids.

After transfecting the MG and helper plasmids into BSR-T7/5 cells, both the viral replicon unit and the control unit are transcribed by the T7 RNA polymerase produced by the BSR-T7/5 cells. The eGFP RNA is then translated by host translation machinery in an IRES-dependent translation manner. For the viral replicon unit, the antiminigenomic RNA generated by the T7 RNA polymerase is coated by N proteins expressed from a helper plasmid (Fig. 1C) to form a ribonucleoprotein and act as replication template (Fig. 1B). The L protein, expressed from a separate helper plasmid (Fig. 1C), will then recognize the promoters located in the leader and trailer sequences of the antigenomic RNA and replicate to produce minigenomic RNA. Minigenomic RNA can also act as a template for antiminigenomic RNA synthesis. Viral polymerase complex proteins also recognize the gene start (GS) signal in the N gene 5’ UTR and gene end (GE) signal in the N gene 3’ UTR of minigenomic RNA and initiate mCherry mRNA transcription. The L protein then modifies mCherry mRNAs to add the 5’ cap and 3’ polyA tail, and finally host translation machinery produces the mCherry protein. Consequently, green fluorescence signal of eGFP and red fluorescence signal of mCherry protein are readouts of this system.

To initially test and validate the minigenomes, all four plasmids (MG, N, P and L) were transfected into BSR-T7/5 cells (MG: 875 ng; N: 312 ng; P: 200 ng; L: 100 ng). mCherry signal was observed in about 9.1% of cells for NiV and 2.5% for HeV transfection group at 72 h post transfection (hpt) (Fig. 2A and B). No mCherry signal was observed in control group which was transfected with the MG, N, and P plasmids without the L plasmid. These data demonstrate that the dual reporter genes minigenome systems for NiV and HeV is functional in BSR-T7/5 cells.

**Fig 2.**
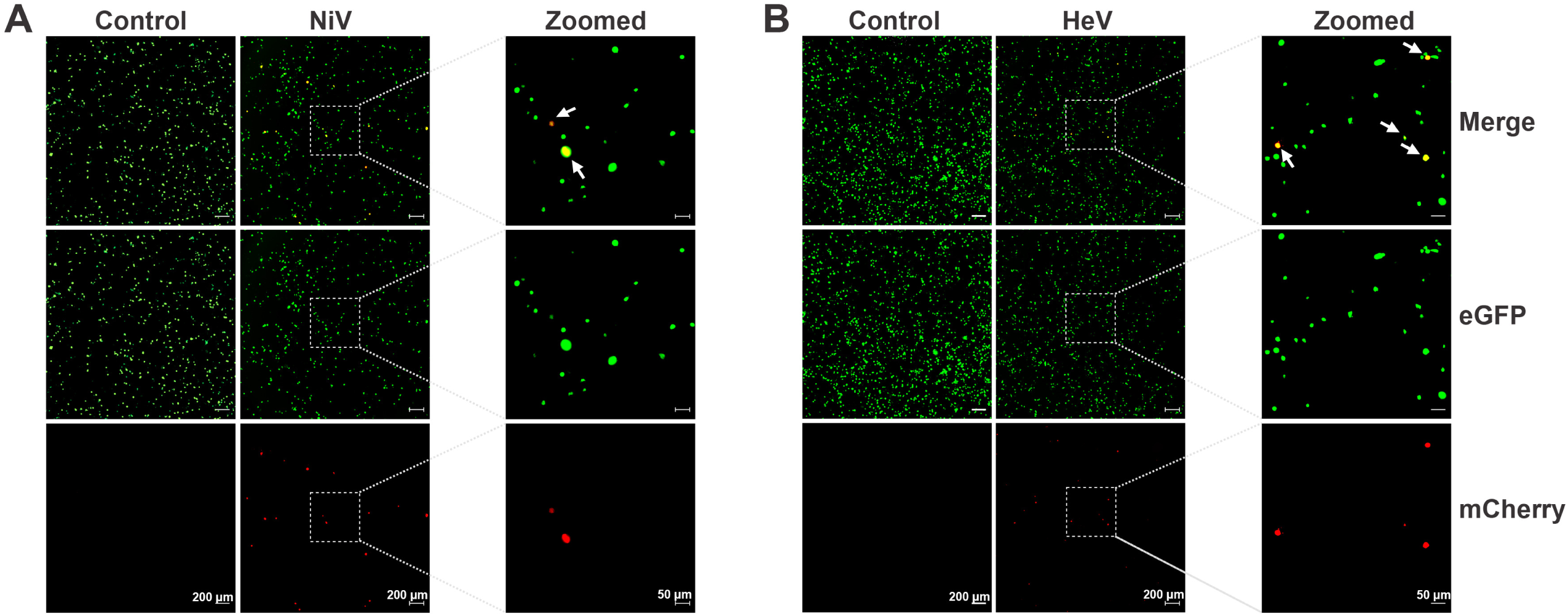
Functional validation of Henipavirus minigenome systems. (**A**) Verification of the NiV Bangladesh strain minigenome system. BSR-T7/5 cells were transfected with NiV MG and three helper plasmids (MG: 625 ng; N: 312 ng; P: 200 ng; L: 100 ng). Control group was transfected with NiV MG, pTM-N, pTM-P of NiV and pTM-1 vector (MG: 625 ng; N: 312 ng; P: 200 ng; pTM1: 100 ng). (**B**) Verification of the HeV Redlands strain minigenome system. BSR-T7/5 cells were transfected with HeV MG and three helper plasmids (MG: 625 ng; N: 312 ng; P: 200 ng; L: 100 ng). Control group was transfected with HeV MG, pTM-N, pTM-P of HeV and pTM-1 vector (MG: 625 ng; N: 312 ng; P: 200 ng; pTM1: 100 ng). Expression of eGFP (Green) and mCherry (Red) was observed at 72 hpt by widefield microscope at 5x magnification. Digital zoomed images are shown in the panel on the right. Cells with both green and red fluorescence signal are indicated by white arrows. Scale bar lengths are indicated.

### Optimization of transfection efficiency of Henipavirus minigenome systems

To optimize the transfection efficiency of both minigenome systems, we tested four different ratios of MG: N: P: L, including ratios that have been previously reported for henipaviruses minigenome system or virus rescue (31–35) (Table 1). “Ratio 2” (MG: 500 ng; N: 150 ng; P: 50 ng; L: 60 ng) resulted in drastically enhanced efficiency at 72 hpt, with 35.6% and 11.0% of cells positive for reporter gene expression for the NiV and HeV minigenomes, respectively, compared to only 2.4% and 6.4% for NiV and HeV respectively in “Ratio 1”, the next best tested (Fig. 3A).

**Table 1.** Transfection ratios of NiV and HeV minigenome system plasmids.

|  | **MG/ng** | **pTM-N/ng** | **pTM-P/ng** | **pTM-L/ng** |
| --- | --- | --- | --- | --- |
| Ratio 1 | 875 | 312 | 200 | 100 |
| Ratio 2 | 500 | 150 | 50 | 60 |
| Ratio 3 | 600 | 600 | 400 | 200 |
| Ratio 4 | 1000 | 410 | 260 | 410 |

**Fig 3.**
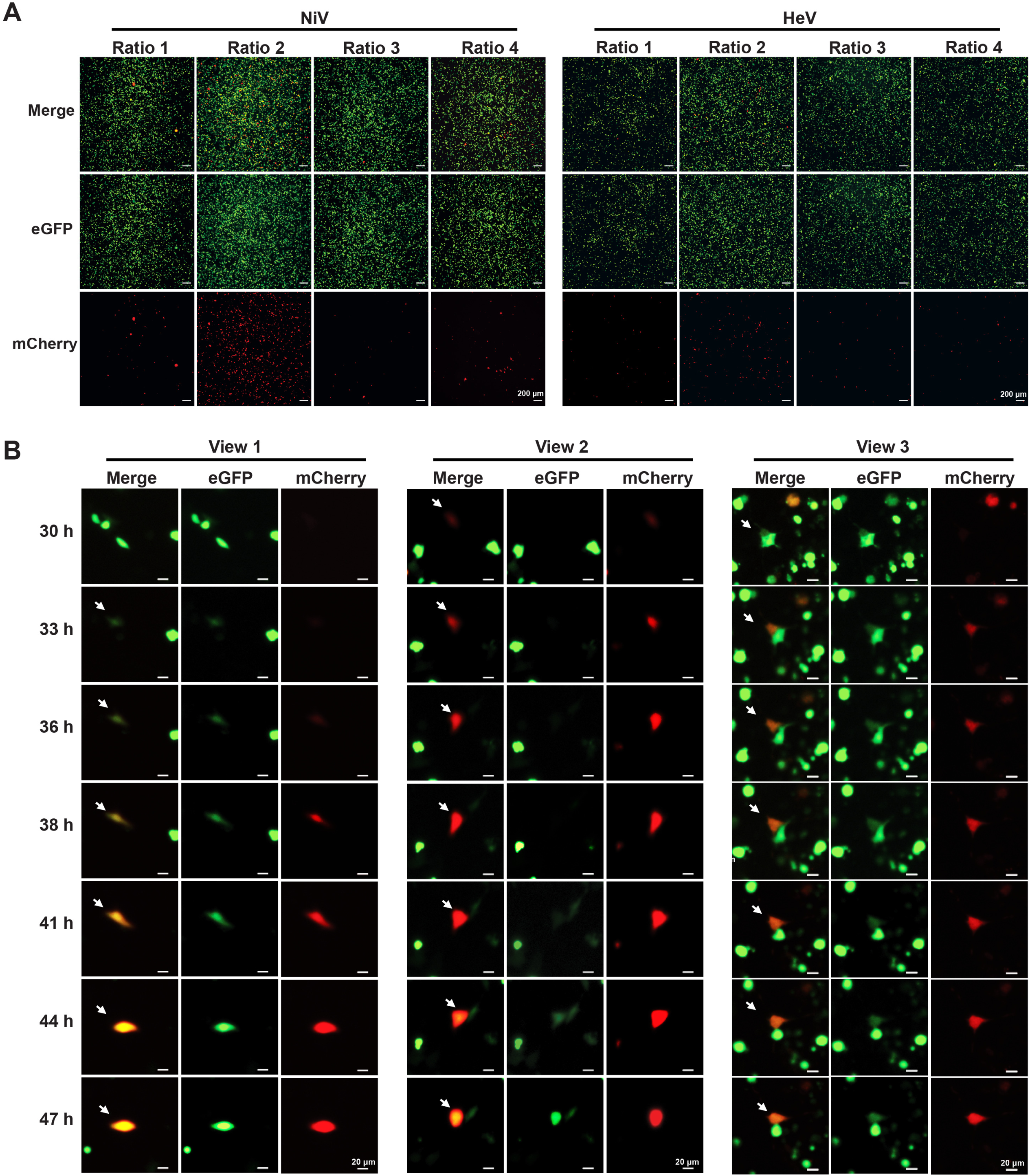
Optimization of Henipavirus minigenome systems. (**A**) BSR-T7/5 cells were transfected with NiV MG alongside the three helper plasmids of NiV (left panel) or HeV MG and the three helper plasmids of HeV (right panel) at 4 different ratios shown in Table 1. Expression of eGFP (Green) and mCherry (red) was observed at 72 hpt. (**B**) Timelapse microscopy images of BSR-T7/5 cells transfected with MG and three helper plasmids of NiV (MG: 500 ng; N: 150 ng; P: 50 ng; L: 60 ng) at 30-47 hpt and eGFP (green) and mCherry (Red) signal are shown at the indicated timepoints. Scale bar lengths are indicated.

Although the “Ratio 2” was identified to have the highest transfection efficiency, we noticed that BSR-T7/5 cells showed several different fluorescence signals among the transfected cells. As expected, most cells showed either only eGFP fluorescence (green) or dual eGFP and mCherry fluorescence (orange), but a few cells showed mCherry only (red), which we did not expect as all mCherry signal should theoretically be accompanied by eGFP in our system (Fig. 1A). We postulated that we were only seeing a snapshot of the reporter protein dynamics and missing the eGFP signal in some cells when the pictures were taken. To test this hypothesis, we used live-cell imaging of cells transfected with the four NiV minigenome system plasmids to investigate the fluorescence signals through time in single cells. Several cells showed bright green fluorescence signal at 33 hpt and then started showing yellow fluorescence signal from 36 hpt gradually increasing expression level of mCherry (Fig. 3B, View 1). In other cases, cells showed single red fluorescence signal at the beginning and turned to orange/yellow at 44 hpt when eGFP was expressed (Fig. 3B, View 2). A different subset of cells showed orange/yellow signal during most of live-cell imaging time course because of similar expression levels of mCherry and eGFP (Fig. 3B, View 3). We did not see any cell that maintained only mCherry signal throughout the entire time-course. These results proved that the fluorescence of single cells was dynamic over time, and we concluded that our minigenome system is working as we expected in its optimized conditions.

### Cross-activity of NiV and HeV polymerase complex proteins

The polymerase complex proteins N, P, and L are essential components for paramyxovirus transcription and replication. As previously discussed, in most cases, homologous support proteins or the full set of heterologous support proteins can replicate of a closely related virus. During optimization of our minigenome systems for NiV and HeV, we assessed the impact of having heterologous components of the polymerase complex in the minigenome efficiency. As previously reported (20), homologous sets of NiV and HeV polymerase components work well in trans to replicate and transcribe the heterologous genome (Fig. 4A and B, Com. 7). However, to our surprise, efficient trans polymerase activity could be seen with all combinations of different polymerase components. As shown by mCherry expression, the NiV MG successfully replicates when HeV N, P, and/or L helper plasmids are used in any combination with the NiV helper plasmids (Fig. 4A). Similarly, all combinations of heterologous NiV helper plasmids work to replicate the HeV MG (Fig. 4B). To test if the observed promiscuity of the henipaviruses polymerase activities was limited to closely related viruses, we used helper plasmids from the paramyxovirus SeV with the NiV minigenome. Transfection of the NiV MG with SeV helper plasmids did not result in mCherry signal, nor could NiV support proteins initiate replication of the SeV minigenome (Fig, 4C). These data suggest that polymerase-associated proteins of NiV and HeV work well with each other but not with other paramyxoviruses.

**Fig 4.**
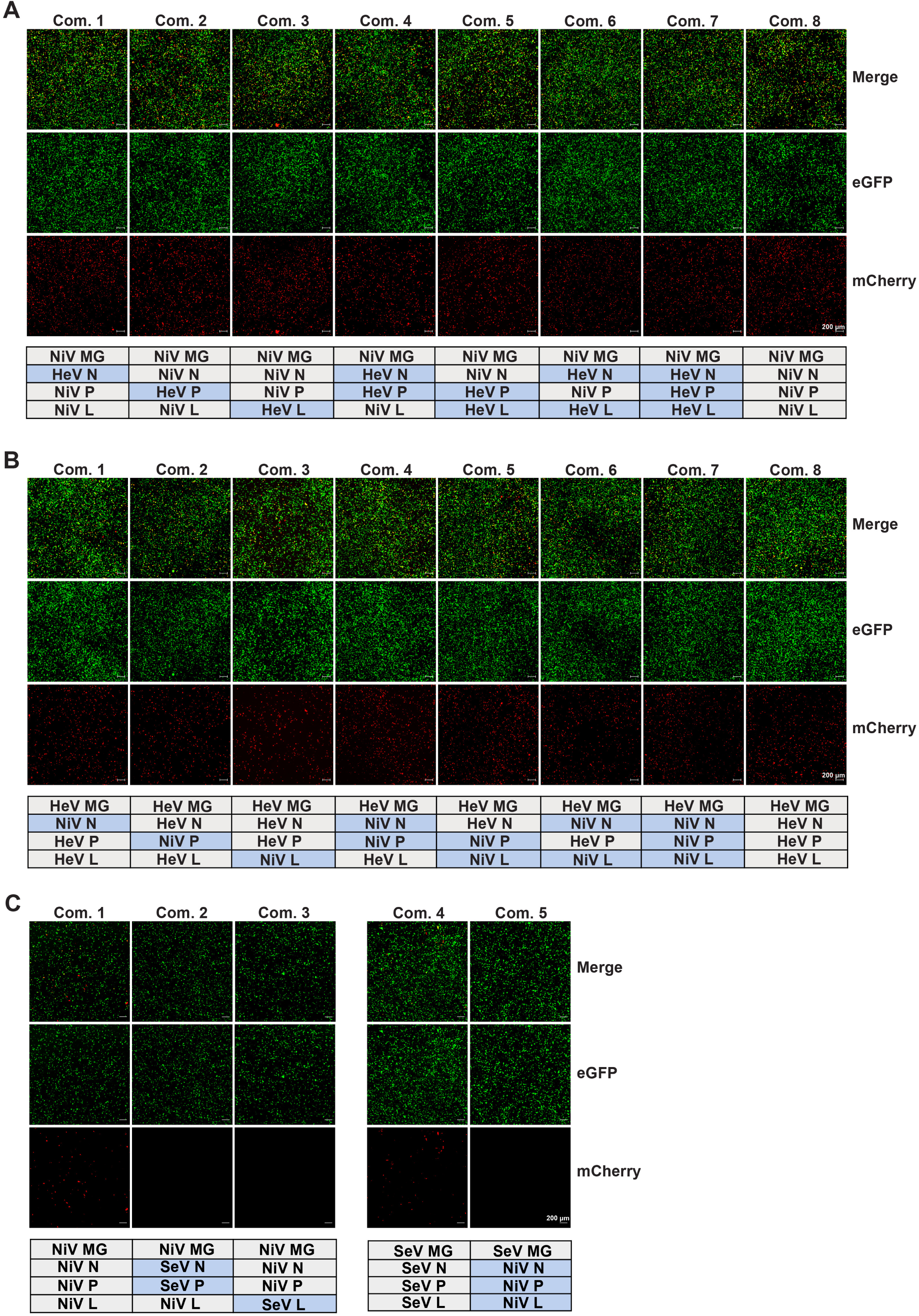
Cross-activity of NiV, HeV, and SeV polymerase complex proteins. (**A**) Cross-activity of different combinations NiV and HeV support proteins in the NiV minigenome system. NiV MG (500 ng) was transfected into BSR-T7/5 cells with the indicated combinations of NiV and HeV helper plasmids (N: 150 ng; P: 50 ng; L: 60 ng). eGFP (Green) and mCherry (red) was observed at 72 hpt. (**B**) Cross-activity of different sets NiV and HeV support proteins in the HeV minigenome system. HeV MG (500 ng) was transfected into BSR-T7/5 cells with different combinations of NiV and HeV helper plasmids (N: 150 ng; P: 50 ng; L: 60 ng). eGFP (Green) and mCherry (red) was observed at 72 hpt. (**C**) Cross-activity verification of NiV and SeV minigenome systems. NiV or SeV MG (500 ng) plasmid were transfected into BSR-T7/5 cells with different combinations of NiV and HeV helper plasmids (N: 150 ng; P: 50 ng; L: 60 ng). eGFP (Green) and mCherry (red) was observed at 48 hpt. Com., Combination. Scale bar shows 200µm.

### Conserved domains for protein-protein interactions likely allow efficient replication, irrespective of support protein combinations

To understand why different combinations of helper plasmids did not affect the replication of Henipavirus minigenomes, as is seen for other *Mononegavirales*, we compared the amino acid sequences of the three polymerase complex proteins of the NiV Bangladesh and HeV Redlands strains used as bases for our minigenomes. We first focused on the N protein since the C-terminal intrinsically disordered domain (N_tail_) of the N protein has 4 defined functional boxes including Box 3, which binds to the C-terminal X domain of viral phosphoprotein (P_XD_) to tether P onto the nucleocapsid template (Fig. 5A) (36, 37). Sequence alignment showed NiV and HeV N proteins share 91.7% identity (defined as the percentage of the same amino acids) and 96.8% similarity (defined as percentage of same amino acids plus conservatively replaced amino acids) (Fig. S1A; Table 2). Only two amino acids (A488I and A492T) within Box 3 showed non-conservative replacement (Fig. 5B). We also compared all amino acid sequences available in the NCBI virus database for NiV and HeV Box 3 and found that most of the Box 3 amino acid sequences of NiV and HeV were conserved (Fig. S2). Differences were mainly shown in 5 novel HeV-g2 variants (UCY33663, UCY33672, UCY33681, UCY33690 and QYC64598) that were isolated between 2013-2020 in Australia (5, 38).

**Fig 5.**
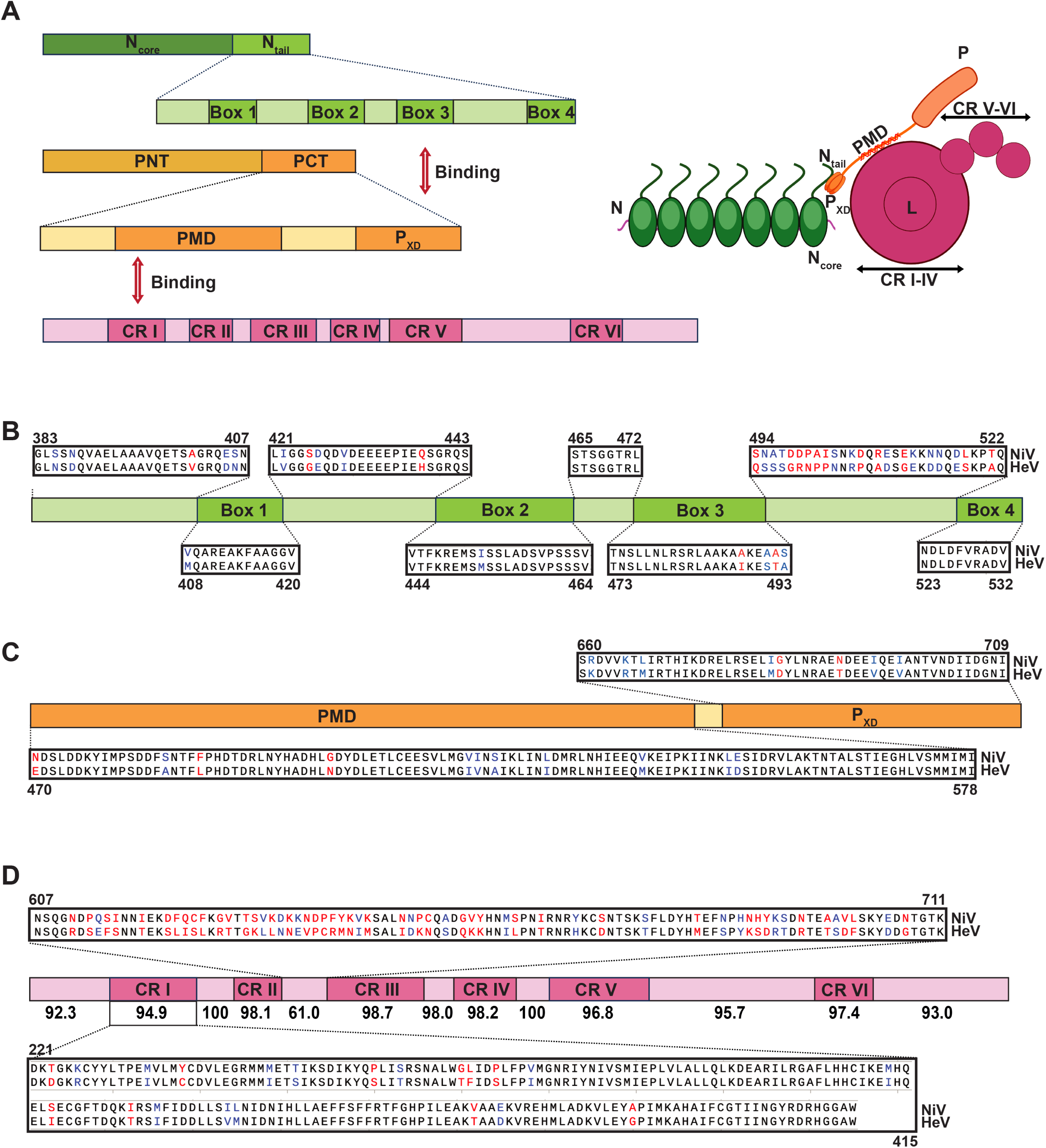
Polymerase complex proteins sequence analysis of NiV Bangladesh and HeV Redlands strains. (**A**) Schematic representation of the interaction between the N and P proteins via key domains. N_core_, structured N-terminal domain. N_tail_, disordered C-terminal domain. PMD, P multimerization domain; PNT, N-terminal region of P; PCT, C-terminal region of P. P_XD_, X domain of P. CR, conserved region. (**B**) Sequence alignment of the N_tail_ of NiV Bangladesh strain and HeV Redlands strain. The first and last amino acids positions for each region based on the NiV Bangladesh strain sequence are displayed above or below. (**C**) Sequence alignment of NiV Bangladesh strain and HeV Redlands strain P protein PMD and P_XD_. PMD and P_XD_ is indicated by orange boxes. The first and last amino acids positions for PMD and P_XD_ are displayed above or below based on the NiV Bangladesh strain sequence. (**D**) Sequence differences analysis of each region of Henipavirus L protein. Schematic representation of Henipavirus L protein, including six conserved regions (CR I-CR VI), which are presented in magenta boxes. Similarities of each region are indicated below the boxes. Sequence alignment of the key CR I and the least conserved region (607–711) between CR II and CR III of NiV Bangladesh strain and HeV Redlands strain are shown individually. The amino acids positions are shown based on the NiV Bangladesh strain sequence. Red text indicates non-similar residues. Blue text represents similar but non-identical residues.

**Table 2.** Identities and similarities of NiV and HeV polymerase complex proteins.

|  | **N protein** | **P protein** | **L protein** |
| --- | --- | --- | --- |
| **Identity** | 91.7% | 66.6% | 87.1% |
| **Similarity** | 96.8% | 77.1% | 94.3% |

The P proteins shared 66.6% identity and 77.1% similarity (Table 2). Although the identity of P protein is lower, sequence alignment result showed most differences were distributed in the N-terminal region of P (PNT) (Fig. S1B). The P_XD_ that binds to N_tail_ was conserved with only two significant amino acids differences (G683D and N690T) (Fig. 5C). Similarly, the P multimerization domain (PMD) that interacts with conserved region I(CRI) of the L protein to recruit L onto the nucleocapsid template (39) was conserved between the two henipaviruses. Alignment of the PMD and P_XD_ sequences of all NiV and HeV strains available from NCBI indicated those two domains were highly conserved, except for one non-similar amino acid substitution in the PMD (G504S) and in P_XD_ (G683D) (Fig. S3 and 4).

Finally, we examined NiV and HeV L protein, a multifunctional enzyme which is conserved among *Mononegavirales*. L has six conserved regions (CRs) named CR I-CR VI (40, 41). CR I, II and III are in the RNA-dependent RNA polymerization (RdRp) domain, CR IV and V are in the cap addition (Cap) domain, and CR VI is in the cap methylation (MT) domain (42). The L proteins of NiV Bangladesh and HeV Redlands shared 87.1% identity and 94.3% similarity (Table 2). The amino acid difference distribution analysis revealed a similarity of over 94.9% for all six CRs. The primary distinction was observed in the region separating CR II to CR III (Fig. 5D), and no significant function has been identified for this area of the protein. We also found that the CR I region of L, which binds primarily to the PMD of the P protein to facilitate P-L interactions, was conserved in all available sequences of henipaviruses, except for the four of the novel HeV-g2 strains (Fig. S5).

These data suggest that dissimilarities within the polymerase complex proteins of NiV and HeV are not concentrated in regions crucial for N, P and L interaction, which may account for the ability of helper plasmids with diverse arrangement combinations to facilitate replication of both NiV and HeV. Furthermore, sequence analysis revealed conservation of these crucial interacting regions across a wide range of NiV and HeV isolates, supporting potential cross-activity between viral proteins.

## DISCUSSION

Henipaviruses are highly pathogenic and global cases are on the rise (5, 43, 44), making the lack of effective therapeutics a pressing concern for human health. The requirement for BSL-4 containment poses significant challenges to the study of these viruses. Minigenome systems are a powerful tool to safely circumvent the need for BSL4 conditions and have been widely used in virus research (45–47), especially for highly pathogenetic agents including Ebola virus, Zika virus, Marburg virus (MARV) and henipaviruses (16, 18, 48, 49). Here, we describe the establishment of a T7 RNA polymerase-based minigenome system for both NiV and HeV and the use of these platforms for analysis of viral polymerase complex proteins cross-activity.

Previously reported minigenome systems for henipaviruses have used chloramphenicol acetyltransferase (CAT), luciferase, or fluorescent proteins such as RFP as the reporter genes to replace all viral structural genes (20, 50, 51). We constructed a bi-cistronic minigenome system including two separate units. The viral replicon unit contained 2 kb of the L gene and was originally designed for further research of henipavirus copy-back viral genomes production during the replication, and the downstream mCherry gene. The control unit express eGFP if the transfection and the T7 polymerase are working well. Unexpectedly, we found single red fluorescent signal in some cells, which has been demonstrated in another bi-cistronic minigenome system for NiV signal based on the relative expression level of eGFP and mCherry protein (50). Using live imaging, we validated the dynamic fluorescent signal based on the relative expression level of eGFP and mCherry protein. We confirmed that all cells eventually express both reporters, although not all cells expressed both reporters at the same time after transfection (Fig. 3B). After optimizing the ratio of four plasmids, we established a robust minigenome system for both NiV and HeV that can be used for further study of henipaviruses. A similar dual reporter strategy could be applied to the research of pro-viral and anti-viral replication factors of henipaviruses and other viruses. Replication and transcription of paramyxoviruses require homotypic support proteins, including N, P and L, although in some condition heterotypic sets are functional among closely related viruses (20, 23, 26, 52). Interestingly, all previous reports on paramyxoviruses show that only support proteins from the same virus can initiate effective replication of a heterologous viral genome or minigenome, highlighting the importance of interactions between N-P and P-L in viral replication (53–55). The viral genome of paramyxoviruses is a negative-sense RNA, which is coated by the N protein to form a ribonucleoprotein (RNP). Functional regions in the N_tail_ bind to P_XD_ to recruit the P protein to the RNP. At the same time, the PMD of the P protein interacts with CR I of the L protein, attaching L to the template. The precise interaction between the polymerase complex proteins is critical for the successful replication of the viral genome and for this reason the polymerase complex has been shown to be a key therapeutic target for paramyxoviruses (52, 53, 56). Consistent with previous reports, we show in our system that the three support proteins of HeV initiate the replication of NiV (20). We also show that N, P, and L of NiV exhibited the ability to replicate HeV minigenome. Strikingly, we also found that various combinations of support proteins enabled replication for both NiV and HeV minigenomes, even when the support proteins were not originating from the same virus (Fig. 4A and B). Interestingly, this phenotype that mixed support proteins from different viruses could initiate the viral replication effectively has not been identified in other paramyxoviruses.

L protein is conserved among paramyxoviruses, while N and P proteins vary. Four boxes exist in N_tail_ of henipavirus N protein and Box 3 binds to P_XD_. This contrasts with both measles virus (MeV) and SeV N_tail_, which have only three boxes and Box 2 interacts with P_XD_ (57–59). The length of P gene varies greatly and is less conserved in paramyxovirus. Even the P gene of two henipaviruses, NiV and HeV, showed a lower identity than N or L (Table 2). Amino acid differences in key regions of polymerase complex proteins may lead to the failure of virus replication with heterologous polymerase complex proteins. This partly explains the inability of the NiV helper plasmids to initiate SeV minigenome replication. The ability of diverse combinations of helper plasmids to facilitate replication of both NiV and HeV minigenomes may be attributed, at least in part, to the conservation of critical protein interaction regions (Box 3 of N, PMD and P_XD_ of P, and CR I of L). Comprehensive sequence alignment suggests potential cross-activity between proteins across a wide range of henipavirus isolates (Fig. S2-5). The increased amino acid variations in emerging HeV-g2 strains underscore the imperative for timely detection and analysis of novel strains. Moreover, these findings suggest that cross-interaction patterns may evolve alongside the emergence of new strains.

While henipavirus outbreaks are currently restricted to Southeast Asia and Australia, the emergence of more henipaviruses and henipa-like viruses raise a serious public health concern of a global pandemic (60, 61). Our findings demonstrate the cross-activity between NiV and HeV, suggesting the possibility of recombinant variants.

It is potential that cross-activity of polymerase complex proteins may not only exist between NiV and HeV, but also happen among different henipaviruses and henipa-like viruses. While this cross-activity may facilitate virus evolution during coinfection of two or more viruses, representing a threat to public health, it also suggests therapeutic potential. Targeting interruption of the interactions of polymerase complex proteins key regions could be pathway to a novel broad antiviral target against henipaviruses.

By utilizing two effective minigenome systems of NiV and HeV, we discovered that different henipavirus polymerase complex proteins have indiscriminate activities and can facilitate heterologous replication of NiV and HeV minigenomes. These data pave the road for future studies on henipaviruses and shed light on our understanding of cross-activity between paramyxovirus polymerase complex proteins, raising novel considerations for viral surveillance and therapeutic development.

## MATERIALS AND METHODS

### Cell culture

BSR-T7/5 cells (hamster kidney cells expressing bacteriophage T7 RNA polymerase, kindly provided by Christopher Basler) were maintained in Dulbecco’s modified Eagle medium (DMEM) (ThermoFisher) supplemented with 10% fetal bovine serum (FBS), L-glutamine 2 mM (Invitrogen), gentamicin 50ng/mL (ThermoFisher), sodium pyruvate 1 mM (Invitrogen) and 400 ug G418 Sulfate (Invitrogen) at 37 °C with 5% CO_2_. Cells were treated with mycoplasma removal agent (MP Biomedicals) before use and screened monthly for mycoplasma contamination with MycoAlert Plus mycoplasma testing kit (Lonza).

### Construction of minigenome system plasmids

To generate the helper plasmids, HeV N, P, L gene sequences were submitted based from the sequence of HeV Redlands strain (GenBank accession No. HM044317) (62) and directly synthesized by IDT (Integrated DNA Technologies). NiV N and L gene sequences were submitted based on the NiV Bangladesh strain sequence (GenBank accession No. AY988601) and synthesized. The NiV P gene was based on NiV Bangladesh genome and internally codon-optimized by IDT to lower the complexity before synthesis. N, P, and L gene open reading frames (ORF) were amplified with PrimeSTAR^®^ Max DNA Polymerase (TAKARA) and ligated between the NcoI and BamHI sites of the pTM1 vector with In-Fusion^®^ Snap Assembly Master Mix (TAKARA) according to the manufacturer’s protocols. SeV Cantell (SeV-C) strain (GenBank accession No. OR764764) (63) helper plasmids were also constructed by cloning and inserting N, P and L gene ORFs into pTM1 vector. All plasmids were confirmed by full length sequencing. For the generation of the dual reporter minigenome plasmids, two separate functional units were constructed separately. Five nucleotides were inserted before the mCherry start codon to ensure the sequence between T7 promotor and T7 terminator of the viral replicon unit conformed the “rule of six”. The minigenome plasmids were constructed by inserting these two units’ sequences into the pSL1180 vector. SeV-C minigenome plasmid was constructed in same way but containing all the viral elements of SeV-C strain.

### Plasmids transfection

BSR-T7/5 cells were seeded in a 24-well plate the day before transfection. Cells were washed twice with PBS (Invitrogen) and incubated in 500 mL Opti-MEM (ThermoFisher) before transfection. BSR-T7/5 cells were transfected with MG, pTM-N, pTM-P and 1 pTM-L using TransIT-LT1 (Mirus) and incubated at 37°C. The negative control group was transfected with MG, pTM-N, pTM-P and pTM1 vector plasmid. Plasmids were transfected into BSR-T7/5 cells according to the ratios showed in Table 1. Fluorescence signal was observed daily after transfection. Images were captured with the 5x and 20x objectives of a Zeiss Axio observer Widefield microscope. Transfection efficiency of viral replicon unit was calculated by mCherry+ cells/eGFP+ cells.

### Time-lapse microscopy

BST-T7/5 cells were observed 30-47 h after transfection with NiV MG and the three helper plasmids of NiV at the “Ratio 2” (Table 1). During observation, cells were maintained in Opti-MEM media at 37°C. eGFP and mCherry fluorescence was visualized using the 5x objectives of a Zeiss Axio observer Widefield fluorescence microscope every 15 mins.

### Sequence analysis

The complete genomes of NiV Bangladesh strain and HeV Redlands strain were obtained from GenBank of the National Center for Biotechnology Information (NCBI). Amino acid identity and similarity of the N, P and L protein amino acids of Bangladesh strain and Redlands strain were analyzed by SnapGene^®^ 6.1.1. Amino acid divergency was represented by the exact match of amino acids (identity) or similarity in amino acid structure (similarity). The complete N, P and L protein sequences of all available Henipavirus isolates submitted on or before Feb 9^th^ 2024 were downloaded from NCBI Virus. Metadata about submitters, counties of isolation, year, host for all sequences were collected from NCBI Virus. Only sequences that represented > 99% of the full-length amino acids sequences of each protein were selected and aligned using MegAlign Pro.

## ACKNOWLEDGEMENTS

The authors acknowledge Roisin Relly for carefully editing the manuscript. This project was funded by a grant from the Novo Nordisk Foundation and the Pandemic Antiviral Discovery (PAD) Initiative (NNF22SA0082041), NIH grant A137062, and the Washington University BJC Investigator program.

